# HDAC6 promote HCCLM3 cell migration and invasion by regulating TGF-β1 pathway

**DOI:** 10.1101/2021.02.23.432488

**Authors:** Zeyu Yu, Xiaoxiao Yang, Guowen Yin

## Abstract

Histone deacetylase 6 (HDAC6) have been demonstrated to play critical roles in the progression of tumor migration and invasion in recent years.However, little research has been done on hepatocellular carcinoma (HCC). This study investigated the role of HDAC6 in the migration and invasion of HCCLM3 cells and its potential mechanism. Western blot and immunohistochemistry were used to detect the expression of HDAC6 in normal hepatic cell line LO2 and HCC cell line HCCLM3. Transforming growth factor 1 (TGF-β1) was proved as a direct target of HDAC6 by Western blot and qRT-PCR.The changes of migration and invasion ability of HCCLM3 cells were confirmed by wound healing and Transwell assays.Morphological changes of HCCLM3 cells were observed by microscope.Our study proved that drownregulated HDAC6 supress HCCLM3 cells metastasis and invasion. Meanwhile, our study verified that HDAC6 regulate EMT by targeting TGF-β1.In conclusion, HDAC6 inhibits the EMT of HCCLM3 cells and further inhibits the migration and invasion of HCCLM3 cells by downregulating the expression of TGF-β1.

## 1. Introduction

Hepatocelluar carcinoma (HCC) is one of the most prevalent tumors worldwide. The frequent metastasisand and recurrence make HCC as a fatal cancer, greatly reduces the 5-year average survival rate of HCC patients. The global incidence of primary liver cancer is the sixth highest among tumors, and the mortality rate has jumped to the second place ^[1]^. Although a variety of treatments have been used to treat liver cancer, However, the overall survival of patients with HCC has not improved significantly over the past two decades, and the mechanisms responsible for the development and progression of HCC remain poorly understood, so the study on the mechanism of its occurrence and development has become particularly important.

In recent years, great progress has been made in the study of cancer. However, due to the absence of specific symptoms in the early stage of liver cancer, patients are often found in the middle and late stage, and there is still no effective treatment for the middle and late stage of liver cancer. Therefore, timely inhibition of metastasis of liver cancer has become the focus of current clinical treatment.Cancer cell metastasis is the difficulty in the treatment of liver cancer and the main reason for the poor prognosis of liver cancer patients^[2]^.Although there are a lot of studies on liver cancer metastasis, the exact molecular mechanism is not yet fully understood.

HDAC6 is a deacetylase that exists in the cytoplasm and regulates a number of important biological processes.It is mainly involved in cell migration, the formation of immune synapses, viral infection and the degradation of allosterins ^[3]^.Studies have shown that HDAC6 can affect tumor development by regulating cell growth, metastasis and apoptosis ^[4–6]^.The purpose of this study was to investigate the effect of HDAC6 on the migration and invasion of human hepatocellular carcinoma HepG2 cells during the development of liver cancer.

## 2. Materials and methods

### 2.1. Cell lines and human tissues

We obtained HCC cells (HCCLM3) and normal hepatic cells (LO2) from Shanghai Institute of Cell Biology, Chinese Academy of Sciences (Shanghai, China). HCC samples (n=9) were acquired from patients undergoing hepatic resection at the Jiangsu Cancer Hospital. Before the collection of specimens, we obtained consent from the candidate patients or their lineal relative. Our study was approved by the Institutional Ethics Board of the Jiangsu Cancer Hospital.

### 2.2. Quantitative real-time PCR

Total RNA was extracted from from HCCLM3 cells and LO2 cells using Trizol reagent (Invitrogen, CA, USA). The total RNA was quantified using NanoDrop 2000 spectrophotometer method (NanoDrop Technologies, MA, USA). Reverse transcription kit was purchased from TaKaRa, Japan.SYBR Green PCR kit was purchased from American Applied Biosystems, RIPA was purchased from Shanghai Yisheng Biological Technology Co., LTD(Shanghai, China).Primers were designed and synthesized by Shenggong Biological Engineering Co., LT D(Shanghai, China). The sequences are as follows:TGF-β1,forward 5’-CTGTAC ATTGACTTCCGCAAG −3’,reverse 5’-TGTCCAGGCTCCAAATGTAG-3’;N-cadherin,forward 5’-CGATAAGGATCAACCCCATACA-3’,reverse 5’-TTCAAAGTCGAT TGGTTTGACC-3’;β-catenin,forward 5’-TGGATTGATTCGAAATCTTGCC-3’,reverse 5’-GAACAAGCA-ACTGAACTAGTCG-3’;Vimentin,forward 5’-ATGTCCACCA GG-TCCGTGT-3’,reverse 5’-TTCTTGAACTCGGTGTTGATGG-3’;GAPDH,forward 5’-TGACTTCAACAGCGACACCCA-3’,reverse 5’-CACCCTGTTGCTGTAGCC AAA-3’. GAPDH was used as internal controls.

### 2.3. Cell transfection and drug treatment

HDAC6 overexpression plasmid (GenePharma Co, Ltd, China) was named P3-HDAC6. Hepatocellular carcinoma cells were transfected with P3-HDAC6 (4 ng) by Lipofectamine 2000 (Invitrogen, CA, USA) when the cells were culture to 80% or 90% confluence. Tubastatin A (MedChemExpress, USA)is a selective inhibitor of HDAC6. SB431542 (MedChemExpress, USA) is a selective inhibitor of TGF-β1. Tubastatin A and SB431542 were dissolvedin Dimethyl sulfoxide(DMSO).They were configured in different concentrations:0μM,10μM,20μM.TGF-β1(Peprotech, USA), which is dissolved in phosphate buffered saline PBS, is a cell growth factor.

### 2.4. Western blotting

The proteins were extracted in RIPA lysis buffer (Yisheng, Shanghai, China) containing 1% protease inhibitor cocktail (Pierce, USA). The NC membranes (VICMED Life Sciences, Xuzhou, China) were incubated overnight at 4 °C with the following specifific primary antibodies: Primary antibodies against HDAC6, TGF-β1,N-cadherin,β-catenin, Vimentin, GAPDH were added for detectionand GA PDH served as an internal control.

### 2.5. Immunohistochemical staining

Tissue slides were deparaffinized and rehydrated.A circle was drawn on the slide around the tissue with a hydrophobic barrier pen and any non-specific binding was blocked. The slides were incubated with the primary antibody (RH DAC6, Abcam, USA) and stained with the chromogen according to a standard staining protocol. Tissue slides were scanned and analyzed by Three-Dimensional Histological Technologies (Pannoramic, 3DHISTECH, Hungary).

### 2.6. Cell morphology

HCCLM3 were plated in six-well plates (2× 10^5^ cells/well) and incubated with TGF-β1(10μg/L).At 24 h post-treatment, the morphology of the cells was examined under an Olympus IX71 microscope (Olympus Corporation, Tokyo, Japan).

### 2.7. Cell migration assay

For the scratch wound-healing motility assay, a scratch was made with a pipette tip when the cells reach the confluence.After being cultured under standard conditions with mitomycin for 48 h, plates were washed twice with fres medi um to remove non-adherent cells and then photographed.The cell migrated from the wound edge were counted and the data were expressed as mean ± SE of three independent experiments.

### 2.8. Cell invasion assay

The transwell inserts (8-μm pores) were filled with 50 μL of a mixture of serum-free DMEM medium and Matrigel.The inserts were then placed in 12-well tissue culture plates containing 10% FBS-medium.After solidification by incubation at 37°C for 4 h,5 × 104 cells in 200 μL medium were placed in upper chambers.Following 48 h of incubation at 37°C with 5% CO2 and in cultur e medium with mitomycin to stop the mitosis, the membranes were fixed with 10% formalin and stained with 0.05% Crystal Violet.The number of cellsthat m igrated through the pores was assessed and the data were expressed as mean ± SE of three independent experiments.

### 2.9. Statistical analysis

Data was presented as means ± SD.Student’s t-test and one-way ANOVA was used for comparisons.All analyses were performed using SPSS software a nd figures were drawn by GraphPad Prism 8.Statistically significance was defin ed as P<0.05 (*),P<0.01 (**).

## 3. Results

### 3.1 Up-regulation of HDAC6 is observed in HCC tissues and cells

To explore the relationship between HDAC6 and the occurrence and development of HCC, we validated HDAC6 expression in 9 paired HCC tissues and their adjacent tissues by way of IHC.We found that the expression of HDAC6 in tumor tissues is distintly higher than that in adjacent non-tumor tissues(Fig. 1A). We then used Western blot with anti-HDAC6 antibody to detect the protein expression level of HDAC6 in the cell lines.The expression of HDAC6 in tumor cell lines was obviously higher than that in normal liver cell lines(*P*<0.05)(Fig. 1B).

**Fig. 1.**
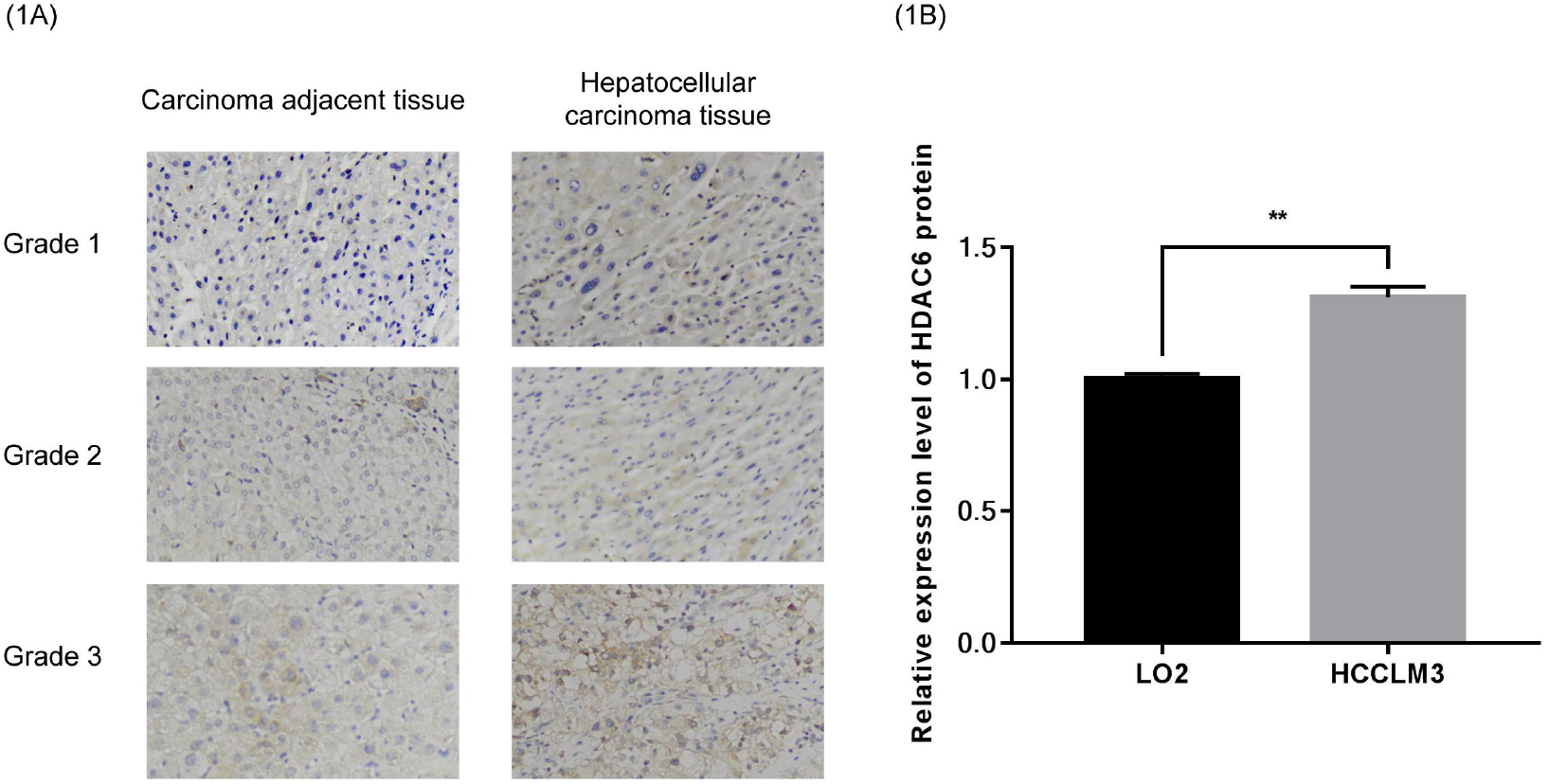
The expression of HDAC6 in HCC tissues and cells. (A) IHC was used to examine the expression of HDAC6 in 9 pairs of HCC tissues and peritumor tissues.(B) HDAC6 protein expression levels were examined in LO2 and HCCLM3 cell line by western blotting analysis (*:*P*<0.05,**:*P*<0.01).

### 3.2 HDAC6 promotes the migration and invasion of HCCLM3

To further explore the potential role of HDAC6 in the development and progression of HCC, we adopted Tubastatin A, it is a selective inhibitor of HDAC6, to supress the expression of HDAC6.Effective silencing of HDAC6 expression in HCCLM3 cells signifificantly decreased the migration ability of HCCLM3 cells as measured in Wound-healing assay(P < 0.01)(Fig.2A). Consistent with the result, the Transwell assays have shown that down-regulating the expression of HDAC6 can suppress the invasion ability of these cells. (Fig.2C-D). Taken together, these results indicated that HDAC6 might play a vital role in promoting the process of HCC.

**Fig. 2.**
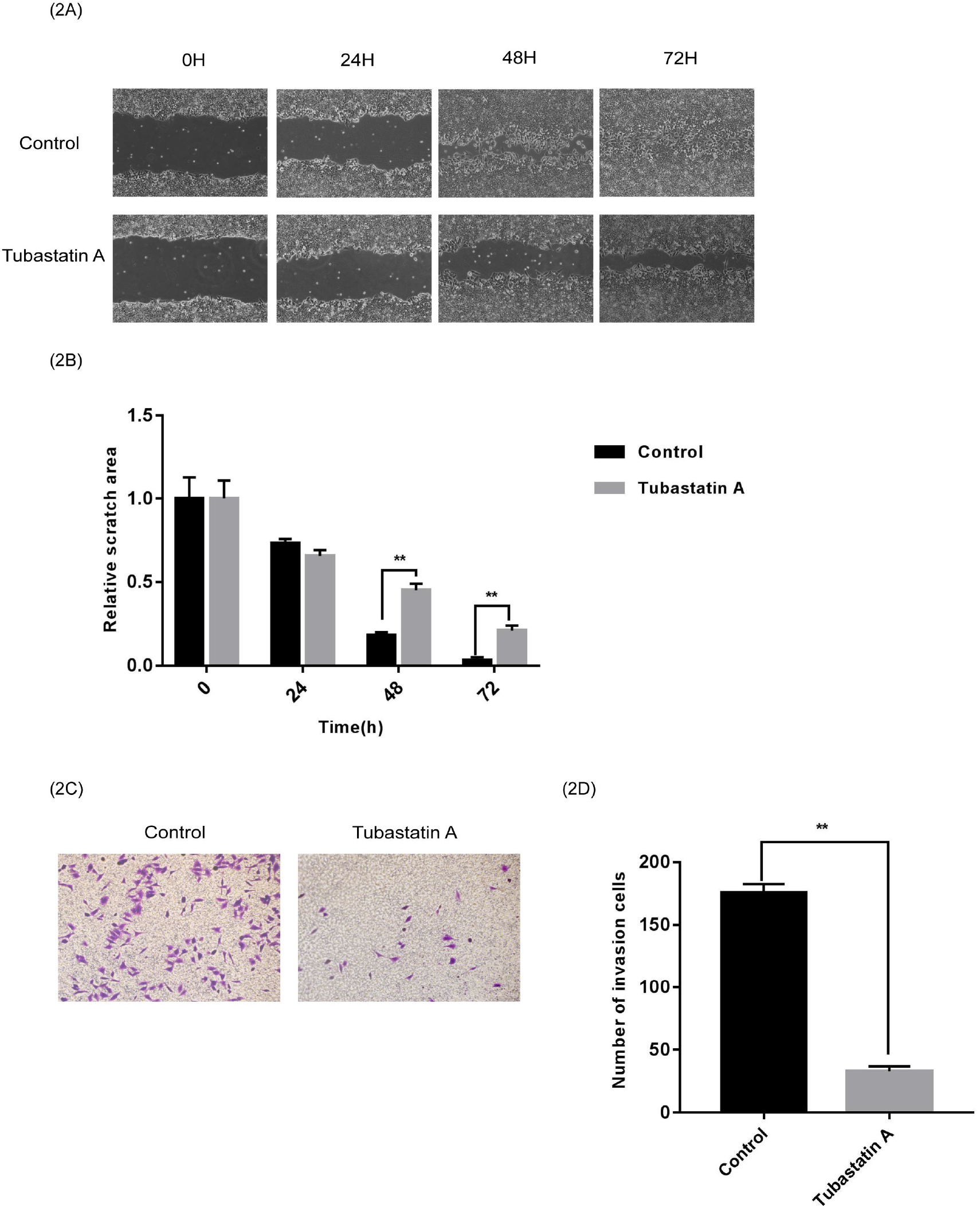
HDAC6 promotes the migration and invasion of HCCLM3. (A-B) Effects of HDAC6 alteration on the migration of the HCCLM3 cells.(C-D) Effects of HDAC6 alteration on the invasion of the HCCLM3 cells.

### 2.3 TGF-β1 can affect the EMT of HCCLM3 cells

Epithelial-mesenchymal transition(EMT) is the process of epithelial cells transformed into mesenchymal cells, which plays an important role in the migration and invasion of tumor cells.EMT phenomenon appears in a variety of malignant tumors. Many researches have shown that TGF-β1 plays an important role in the EMT process.We asked the question whether down-regulation of TGF-β1 could affect EMT in HCCLM3. As shown in(Fig.3A-B),down-regulation of TGF-β1 has a negative infiuence of the expression of these proteins which is crucial in the process of EMT.What is more, down-regulation of TGF-β1 also affect the morphological characteristics of HCCLM3 cells(Fig.3C). Taken together, our results suggest that TGF-β1 might be an important positive regulator of the process of EMT.

**Fig. 3.**
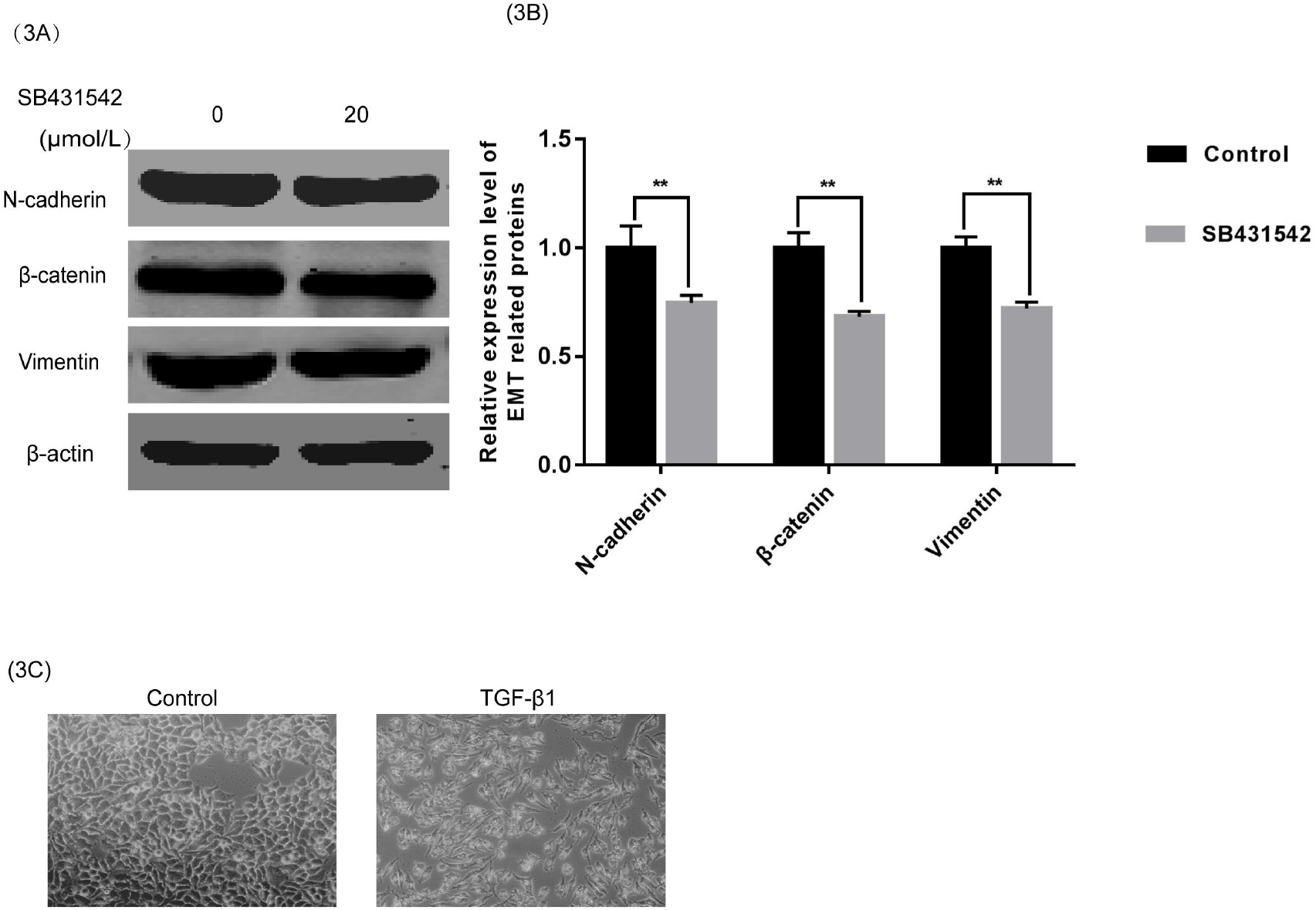
TGF-β1 affect the EMT of HCCLM3 cells. (A-B) The expression of EMT related proteins were detected by WB in the HCCLM3 cells treated with SB431542.(C) Microscopy shows the changes of HCC-LM3 cells morphology induced by TGF-β1.

### 2.4 HDAC6 is required for the process of EMT and the activation of TGF-1

EMT plays a very critical role in the migration and invasion of tumor cells.To understand whether HDAC6 is involved in the process of EMT, we supressed the expression of HDAC6 with Tubastatin A in HCCLM3 cells.Then we find the levels of the proteins and mRNA of the EMT related markers are both down-regualted (Fig.4A-C). These results show that HDAC6 is of great significance to the process of EMT in HCCLM3 cells.Previous studies have shown that TGF-β1 might be an important positive regulator of the process of EMT.We asked the question whether HDAC6 have something to do with TGF-β1.To explore the relationship between HDAC6 and TGF-β1,we have adopted PCR and Western Blot technology.The results showed that supressing HDAC6 could down-regulate the expression of TGF-β1 (Fig.4D-E) and block the EMT process.

**Fig. 4.**
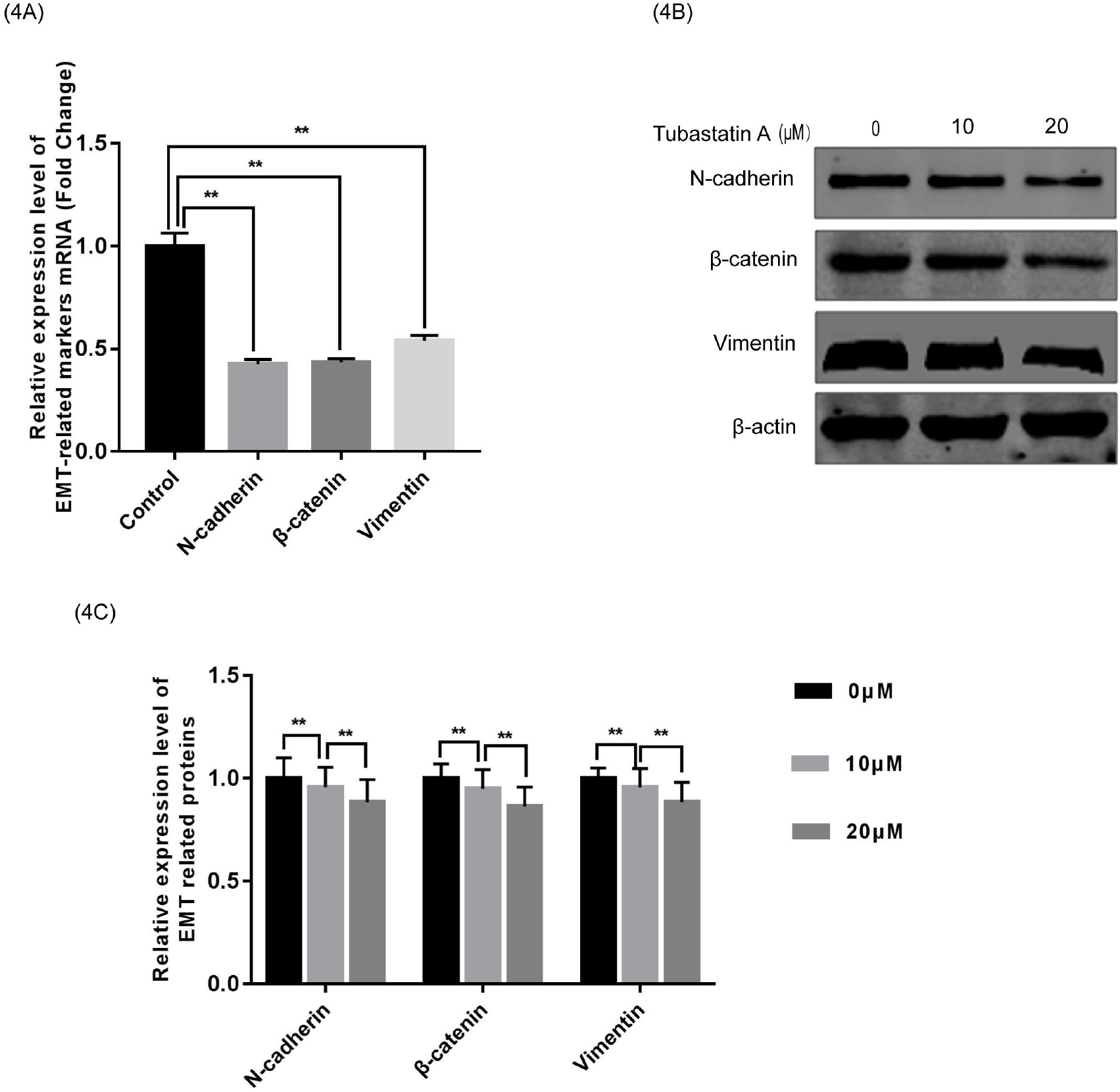
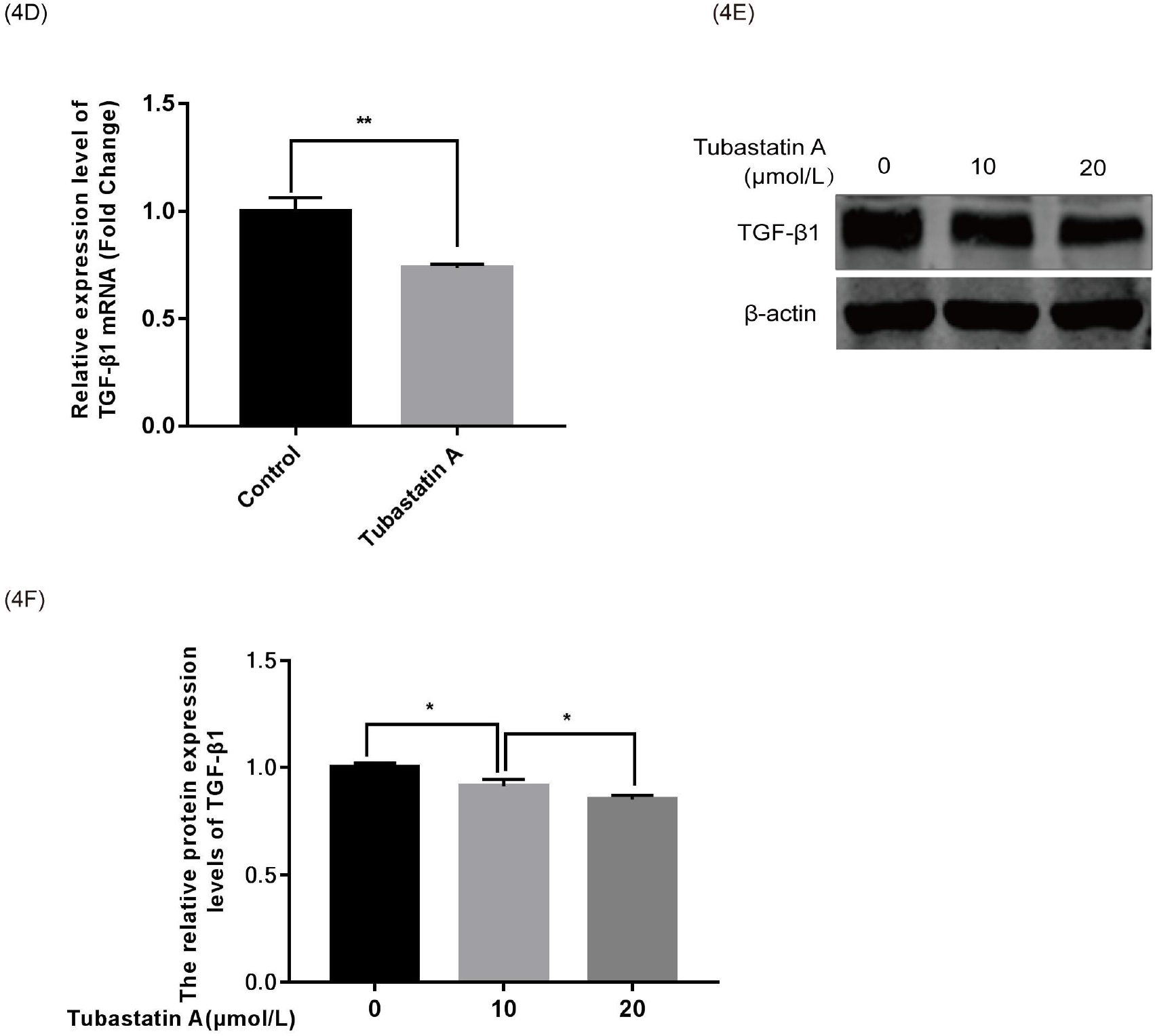
TGF-β1 affect the EMT of HCCLM3 cells. (A-C)Protein and mRNA expression levels of EMT-related markers in HDAC6 low-expressed HCCLM3 cells were detected by WB and qRT-PCR.(D-F) WB and qRT-PCR analysis of the Proteins and mRNA expression levels of EMT-related markers in HDAC6 low-expressed HCCLM3 cells.

### 2.5 HDAC6 activated EMT via TGF-1 expression in HCCLM3 cells

We can draw these conclusions from previous research that the expression of TGF-β1 is influenced by HDAC6 and TGF-β1 can affect the process of EMT. Therefore, we hypothesized that HDAC6 could facilitate the process of EMT though TGF-β1. To test this hypothesis, We overexpressed HDAC6 with plasmids in HCCLM3 cells. Then, we analyzed the expression of EMT related proteins levels in HCCLM3 cells treated with empty drugs or SB431542.As shown in (Fig.5A-B), the results indiacted that HDAC6 might influence the process of EMT though the TGF-β1 pathway, what is consistent with the previous conclusion.

**Fig. 5.**
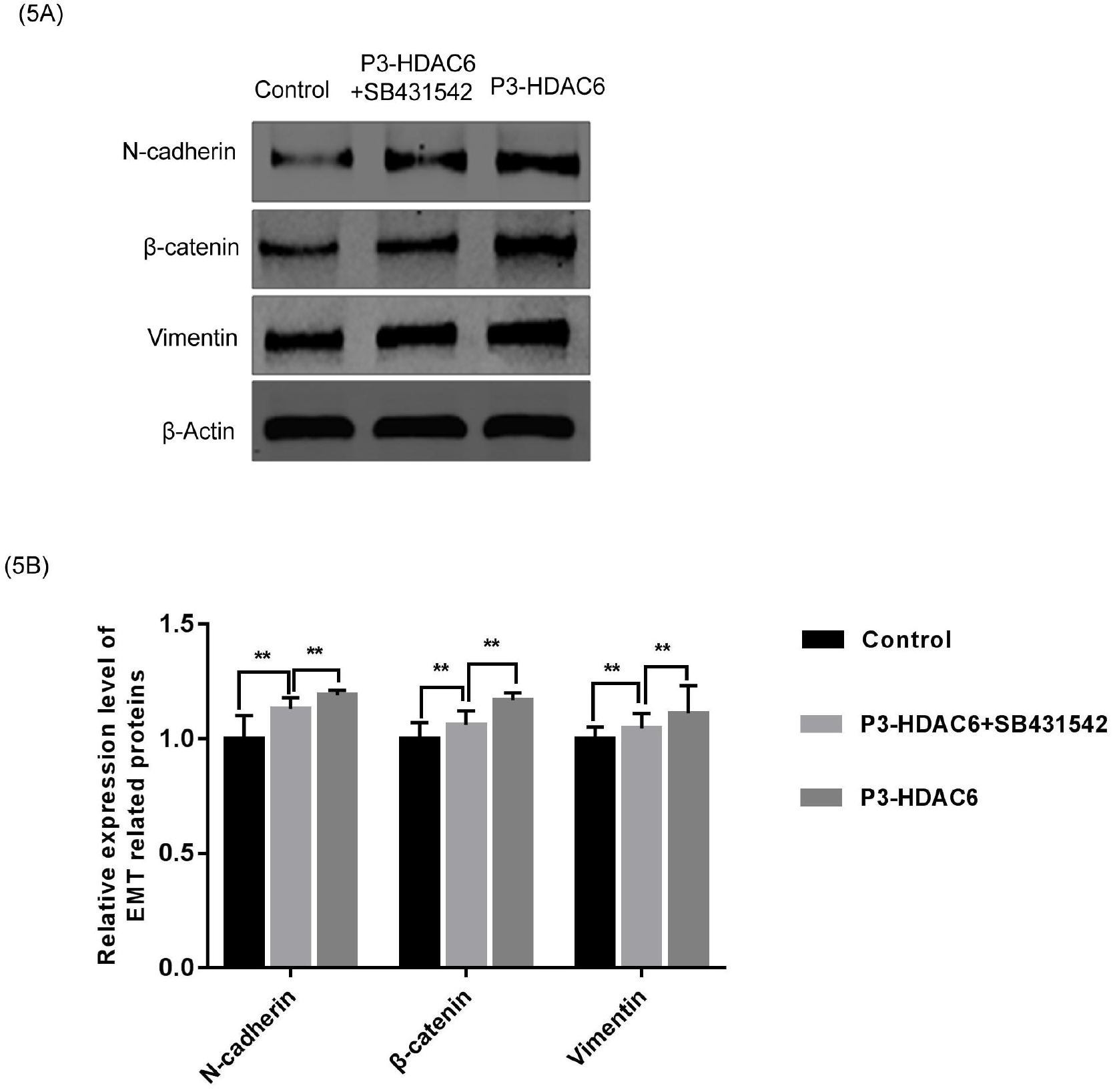
HDAC6 activated EMT via TGF-1 expression in HCCLM3 cells. (A-B)The expression levels of EMT related proteins in the HCCLM3 cells, which are treated with different drugs, were detected by WB.

### 2.6 HDAC6 promotes migration and invasion of HCCLM3 cells by TGF- 1

In order to further understand the role of HDAC6 in the occurrence and development of HCC, we regulated the expression of HDAC6 and TGF-β1, and then observed the changes in the metastasis and invasion ability of HCCLM3 cells.The result of Wound-healing assays showed that up-regulation of HDAC6 could enhance the migration of HCCLM3 cells.However, simultaneous up-regulation of HDAC6 and down-regulation of TGF-β1 would make the migration ability of HCCLM3 cells weaker than the previous one, but stronger than the primary(Fig.6A-B). As expected, we came to a similar conclusion in the Transwell assays(Fig.6C-D).

**Fig. 6.**
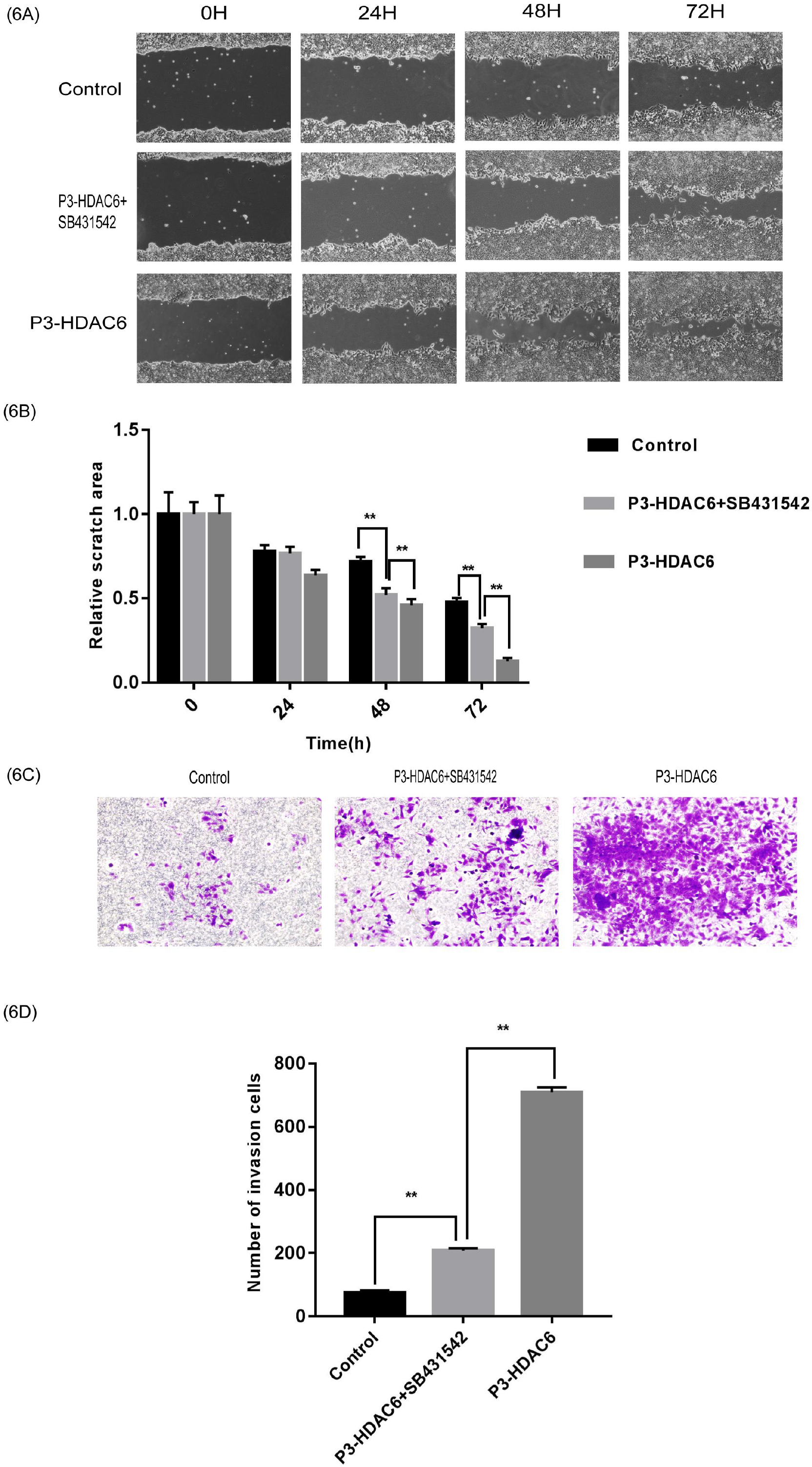
HDAC6 promotes migration and invasion of HCCLM3 cells by TGF- 1. (A-B)The ability of migration in the HCCLM3 cells treated with different drugs was detected by Wound-healing. (C-D)The ability of invasion in the HCCLM3 cells treated with different drugs was detected by Transwell assays.

## 4. Discussion

HDAC6, a member of the histone deacetylase family, inhibits gene transcription through deacetylation of histones, thereby affecting the growth state of cancer cells^[7]^. HDAC6 inhibits not only the transcription of oncogenes but also the transcription of oncogenes, so it can play a completely different role in different cancer cells. The role of HDAC6 in hepatocellular carcinoma (HCC) is still controversial. Based on previous studies, it is found that studies come to conflicting conclusions are often used different HCC cell lines, and there is no totally opposite conclusion in studies with the same HCC cell line. These studies indicate that the effect of HDAC6 on HCC is quite different in different HCC cells^[7–12]^. Now we’re looking at two possibilities.Firstly, in different hepatocellular carcinoma cell lines, due to the different intracellular environment of different hepatocellular carcinoma cells, the microstructure of HDAC6 may be changed, leading to the change of its function. Secondly, since gene methylation and histone deacetylation synergistic inhibit gene transcription, and histone deacetylation inhibiting gene transcription capacity depends on the number of DNA methylation CpG sites, When the number of DNA methylation CpG sites reaches a certain number, the transcriptional inhibition of genes mainly depends on DNA methylation^[13]^.However, when the number of CpG sites for DNA methylation was low, the ability of histone deacetylation to inhibit gene transcription was dominant^[14]^. In different liver cancer cell lines, the number of methylation CpG sites in the promoter, enhancer, silencer and other cis elements of the target gene may be different, so the deacetylation function of HDAC6 has different effects on the transcription of the target gene, and the function of HDAC6 has a series of differences in different liver cancer cells.

EMT is the transformation of epithelial cells into mesenchymal cells, which gives cells the ability to metastasize and invade.EMT phenomenon appears in a variety of malignant tumors, and it is an important reason for the occurrence and development of malignant tumors^[15]^. The results of this experiment suggest that HDAC6 can promote the migration and invasion of cancer cells by promoting the EMT of HCCLM3 cells.

TGF-β1 is involved in the regulation of cell migration, adhesion and apoptosis, and it plays an important role in the development of tumors^[16]^. TGF-β1 pathway is considered to be the most influential signaling pathway involved in the EMT process, and EMT plays an important role in the development and metastasis of HCC^[17–18]^. We found that the expression of TGF-β1 was positively correlated with the expression of HDAC6 in liver cancer cells. With the decrease of HDAC6, the expression of TGF-β1 decreased correspondingly.

In summary, we demonstrated that HDAC6 promoted the EMT of HCCLM3 cells through the TGF- TGF 1 pathway, thereby promoting the migration and invasion of HCCLM3 cells. These results are important for the exploration of the mechanism of liver cancer cell metastasis.

